# Opportunities for risk-taking during play alters cognitive performance and prefrontal inhibitory signalling in rats of both sexes

**DOI:** 10.1101/2023.08.29.555263

**Authors:** Ate Bijlsma, Evelien E. Birza, Tara C. Pimentel, Janneke P.M. Maranus, Marieke J.J.M van Gaans, José G. Lozeman-van t Klooster, Annemarie J.M. Baars, E.J. Marijke Achterberg, Heidi M.B. Lesscher, Corette J. Wierenga, Louk J.M.J. Vanderschuren

**Author notes:** **Correspondence:** Corette Wierenga, Louk Vanderschuren.

## Abstract

Social play behaviour is a rewarding activity that can entail risks, thus allowing young individuals to test the limits of their capacities and to train their cognitive and emotional adaptability to challenges. Here, we tested in rats how opportunities for risk-taking during play affect the development of cognitive and emotional capacities and medial prefrontal cortex (mPFC) function, a brain structure important for risk-based decision-making. Male and female rats were housed socially or social play-deprived (SPD) between postnatal day (P)21 and P42. During this period, half of both groups were daily exposed to a high-risk play environment. Around P85, all rats were tested for cognitive performance and emotional behaviour after which inhibitory currents were recorded in layer 5 pyramidal neurons in mPFC slices. We show that playing in a high-risk environment altered cognitive flexibility in both sexes, and improved behavioural inhibition in males. High-risk play altered anxiety-like behaviour in the elevated plus maze in males and in the open field in females, respectively. SPD affected cognitive flexibility in both sexes and decreased anxiety-like behaviour in the elevated plus maze in females. We found that synaptic inhibitory currents in the mPFC were increased in male, but not female, rats after high-risk play, while SPD lowered PFC synaptic inhibition in both sexes. Together, our data show that exposure to risks during play affects the development of cognition, emotional behaviour and inhibition in the mPFC. Furthermore, our study suggests that the opportunity to take risks during play cannot substitute for social play behaviour.

## Introduction

Play behaviour is an intrinsically rewarding activity that is abundant among the young of humans and most other mammalian species (Pellis and Pellis, 2009; Graham and Burghardt, 2010). It is typically expressed under safe circumstances in which individuals can experiment with their behaviour and develop motor, social, cognitive and emotional skills necessary for optimal functioning later in life (Špinka et al., 2001; Pellis and Pellis, 2009; Gray, 2017; Nijhof et al., 2018; Sgro and Mychasiuk, 2020). Importantly, when exploring and experimenting with behaviour, playful situations may entail risks, such as the risk of rejection, injury or failure. Although the term “risk” often has negative connotations, risky activities provide ample opportunities to acquire new skills, explore one’s physical boundaries (Brussoni et al., 2012; Lavrysen et al., 2017) and learn to cope with anxiety through exposure to fearful stimuli (Allen and Rapee, 2005; Sandseter and Kennair, 2011). Case studies, for example, suggest that play in a natural environment can increase children’s problem-solving skills and the ability to successfully cope with difficult or challenging life experiences (McArdle et al., 2013), while limited risky and outdoor play have been associated with lower self-esteem and academic achievements (Tremblay, 2011; Tremblay et al., 2015). There is substantial concern that opportunities for children to engage in free and risky play have declined over the past decades in Western societies (Valentine and McKendrick, 1997; Weir et al., 2006; Gray, 2011; Brussoni et al., 2012; Little et al., 2012). Although it is generally assumed that risk-taking during play is important to prepare the individual for challenges later in life, little empirical research has been done to explore the contribution of risks to the development of brain and behaviour.

Animal studies, especially in rats, have been valuable to investigate the developmental functions of social play behaviour (Vanderschuren and Trezza, 2014; Pellis et al., 2023). Rats that had been deprived of social play between weaning and adolescence, a period in which social play is especially abundant, display inappropriate behaviour in a social conflict situation (Van Den Berg et al., 1999; Von Frijtag et al., 2002), altered impulse control and decision making (Baarendse et al., 2013; Bijlsma et al., 2022) and increased sensitivity to substances of abuse (Whitaker et al., 2013; Baarendse et al., 2014; Lesscher et al., 2015). At the neuronal level, altered functioning of the prefrontal cortex (PFC) (Bell et al., 2009; Wall et al., 2012; Baarendse et al., 2013; Bijlsma et al., 2022, 2023), a brain structure that is important for higher cognitive, so-called executive functions (e.g. working memory, impulse control, attention, planning and decision making), has been observed in play-deprived animals. Indeed, we have recently demonstrated that deprivation of social play affects the synaptic connectivity in the medial prefrontal cortex (mPFC) of adult rats; we found a reduction in specific inhibitory synaptic inputs onto layer 5 pyramidal cells (Bijlsma et al., 2022, 2023). These findings support the notion that social play behaviour subserves the development of emotional and cognitive capacities and their PFC substrate. However, it remains unclear how the opportunity to take risks during play affects the development of emotion, cognition and PFC function.

Sandseter (2007) has described six categories of “risky” play in children: 1) play with great heights; 2) play with high speed; 3) play with harmful tools; 4) play near dangerous elements; 5) rough-and-tumble play; and 6) play where children can ‘disappear’/get lost (Sandseter, 2007). Currently, social play behaviour assessment in rats entails only one of these categories of risky play, i.e. rough-and-tumble play, but not the other ones. Moreover, the typical laboratory housing settings for rodents are limited in the actual risks that the animals may encounter. Here we therefore combined social play deprivation with opportunities for risk-taking in play to create four rearing conditions in rats that reflect a continuum of play behaviour, ranging from no play to a combination of social and risky play. These conditions were used to answer the following questions: 1) Does the opportunity for risk-taking behaviour during play alter cognitive flexibility, behavioural inhibition and anxiety-like behaviour?; 2) How does the opportunity for social play affect cognitive flexibility, behavioural inhibition and anxiety-like behaviour?; 3) How does play manipulation affect the maturation of the mPFC?; and 4) Do the consequences of high-risk play and social play deprivation differ between males and females?

## Materials & methods

### Animals and housing conditions

Three batches of 48 male and one batch of 48 female Lister Hooded rats were obtained from Charles River (Sulzfeld, Germany). They arrived on postnatal day (P)14 in litters of eight with nursing mothers. All rats were subjected to a reversed 12:12h light-dark cycle (lights on 07:00, lights off 19:00) with ad libitum access to water and food. All experiments (Fig. 1A) were conducted during the active phase of the animals (9:00 - 17:00). Rats were weighed and handled at least once a week throughout the experiment. One week before the start of behavioural testing, the rats were subjected to a restricted diet of 4.5 grams of chow per 100 grams of body weight. In this way, animals were kept at 90% of their body weight for the duration of behavioural testing. Body weights did not differ significantly between the three male batches. Rats were provided with 30 sucrose pellets (45mg, BioServ) in their home cage before their first exposure to the pellets in the operant conditioning chamber to reduce potential food neophobia. The age at the start of behavioural experimentation was comparable for each task and batch. All experiments were approved by the Animal Ethics Committee of the Utrecht University and the Dutch Central Animal Testing Committee. They were conducted in agreement with Dutch (Wet op de Dierproeven, 1996; Herziene Wet op de Dierproeven, 2014) and European legislation (Guideline 86/609/EEC; Directive 2010/63/EU).

**Figure 1.**
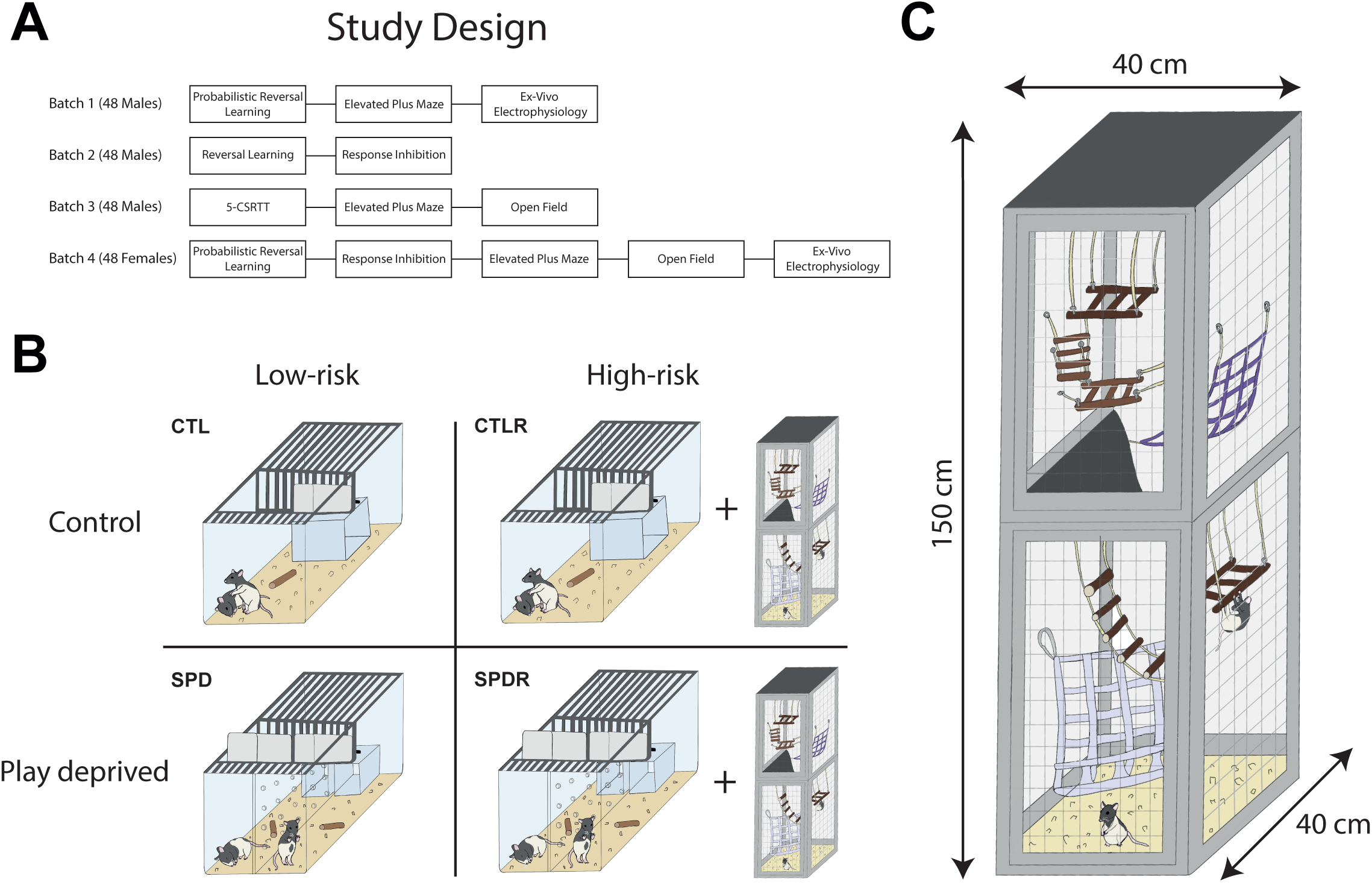
Study design. (A) Study overview. (B) Combination of a social play deprivation paradigm and high-risk play enrichment resulting in four rearing conditions that reflect a continuum of play behaviour: control (CTL-LR), social play deprivation (SPD-LR), control risky play (CTL-HR) or social play deprivation risky play (SPD-HR). (C) The risky play cage was enriched with multiple ladders, plateaus and other objects to interact with.

### Social play deprivation and risky play

Rats were weaned on P21 and were either subjected to the control (CTL-LR), social play deprivation (SPD-LR), control risky play (CTL-HR) or social play deprivation risky play (SPD-HR) group, resulting in 4 groups of 12 rats (Fig. 1B). All rats were housed in pairs with a littermate during the entire experiment. From P21 to P42, a transparent plexiglass divider containing small holes was placed in the middle of the home cage of SPD-LR and SPD-HR rats creating two separate but identical compartments. Social play-deprived rats were therefore able to receive visual, olfactory and auditive cues from one another. The holes in the plexiglass allowed the rats to have limited physical interaction but no opportunity to physically engage in play. The divider was removed on P42 and SPD-LR/SPD-HR rats were housed in the same pairs for the remainder of the experiment. CTL-HR and SPD-HR were housed under the same conditions as the CTL-LR and SPD-LR groups but were transferred to a “risky play cage” twice a day for 30 minutes during the deprivation period (P21 – P42). The risky play cage measured 150 x 40 x 40 cm (H x W x D) and was enriched with multiple ladders, plateaus and other objects to interact with (Fig. 1C). CTL-HR rats were placed together in the risky play cage, while SPD-HR rats were alone. The risky play cages were designed to allow the rats more opportunities to take risks during play, in comparison to their normal home cage.

### Probabilistic reversal learning (PRL) task

#### Apparatus

Behavioural testing was conducted in operant conditioning chambers (Med Associates, Georgia, USA) enclosed in sound-attenuating cubicles equipped with a ventilation fan. Two retractable levers were located on either side of a central food magazine into which sugar pellets could be delivered via a dispenser. A LED cue light was located above each retractable lever. A white house light was mounted in the top centre of the wall opposite the levers. Online control of the apparatus and data collection was performed using MED-PC (Med Associates) software.

#### Pre-training

Rats were first habituated once to the operant chamber for 30 min in which the house light was illuminated and 50 sucrose pellets were randomly delivered into the magazine with an average interval of 15 s between reward deliveries.

#### Phase 1

On the subsequent days, the rats were trained for 30 min under a Fixed-Ratio 1 (FR1) schedule of reinforcement for a minimum of two consecutive daily sessions. A FR1 session started with the illumination of the house light and the insertion of both levers, which remained inserted for the remainder of the session. A response on one of the levers resulted in the delivery of a sucrose pellet into the magazine. There was no limit other than time (max 30 min) or the number of times a rat could press the levers (max 100). To proceed to phase 2, the rat had to obtain an average of at least 50 rewards over two completed sessions. In case a rat obtained a lower number of rewards during the first two sessions, it was further trained on subsequent days until the criterion was met.

#### Phase 2

A trial started with the presentation of the left lever, the right lever, or both levers and pressing either lever was reinforced under a FR-1 schedule. In this phase of training, the levers were only retracted after a response, and the animals were subjected to the same sequence of events as during a reinforced trial in the probabilistic reversal learning task. When all animals made at least 100 responses in a session during this phase they progressed as a group to the next phase.

#### Phase 3

Rats were familiarized with the probabilistic nature of the PRL task, in which both levers were presented and a lever press resulted in 80% reward delivery instead of 100%. Levers were presented until pressed. Rats were trained to a criterion of at least 50 rewards and to make at least one lever press in more than 80% of the trials before progressing to the probabilistic reversal learning phase; this required ∼3-4 days.

#### Probabilistic reversal learning

The protocol used for this task was modified from those of previous studies (Bari et al., 2010; Dalton et al., 2016; Verharen et al., 2020; Bijlsma et al., 2022). At the start of each session, one of the two levers was randomly selected to be ‘correct’ and the other ‘incorrect’. A response on the ‘correct’ lever resulted in the delivery of a reward on 80% of the trials, whereas a response on the ‘incorrect’ lever was reinforced on 20% of trials. Each trial started with a 5 s ITI, followed by the illumination of the house light and the insertion of both levers into the chamber. After a ‘correct’ response, both levers retracted. In case the rat was rewarded, the house light remained illuminated, whereas the house light extinguished in case the rat was not rewarded on the ‘correct’ lever. An ‘incorrect’ response or a failure to respond within 30 s after lever insertion (i.e. omission) lead to the retraction of both levers and extinction of the house light so that the chamber returned to the ITI state. When the rat made a string of 8 consecutive trials on the ‘correct’ lever (regardless of whether they were rewarded or not), contingencies were reversed, meaning that the ‘correct’ lever became the ‘incorrect’ lever and the previously ‘incorrect’ lever became the ‘correct’ lever. This pattern repeated throughout a daily session. Daily sessions were completed upon performing 200 (males) or 140 trials (females). Female rats were subjected to a lower maximum of trials as they became sated faster which lead to a lower response rate in later trials. The possible number of reversals made was solely limited by the number of trials in a session.

#### Trial-by-trial analysis

This analysis was performed to assess the shifts in choice behaviour between subsequent trials and to investigate the sensitivity to positive and negative feedback. Depending on whether the rat received a reward or not, it can press the same lever on the subsequent trial or shift towards the other lever, resulting in 4 different possibilities:

- Win-stay: Same lever press on the subsequent trial after receiving a reward.
- Win-shift: Opposite lever press on the subsequent trial after receiving a reward.
- Lose-stay: Same lever press on the subsequent trial after receiving no reward.
- Lose-shift: Opposite lever press on the subsequent trial after receiving no reward.

### Response inhibition (RI) task

#### Apparatus

Behavioural testing was performed in the operant chambers as used for the PRL, except that they were equipped with a shock grid floor and a tone generator (4500 Hz).

#### Pre-training

Rats were first habituated for four days to the operant chamber for 30 minutes during which the house light was illuminated. Fifty sucrose pellets were randomly delivered into the magazine with an average of 15 s between reward deliveries. This phase was followed by a training phase in which, after an initial 20 s before the start of the first trial, rats had 40 s per trial to retrieve a sucrose pellet. When the sucrose pellet was not retrieved, a new trial started but no new reward was given. The session ended after 60 trials.

#### Response inhibition

The protocol used for this task was modified from Verharen et al. (2019) (Verharen et al., 2019). In contrast to the study by Verharen et al., in which the shock intensity was adjusted for each individual, we used the same shock intensity for every rat. A response inhibition (RI) session consisted of 60 trials of 40 seconds of which 30 were non-stimulus (NS) trials and 30 were shock trials. In NS trials, rats were allowed to retrieve the sucrose reward directly after its delivery. During the first 12 seconds of shock trials, two cue lights were turned on and a tone was produced by a speaker in the top right corner on the same wall as the house light. When rats collected the reward during the 12 seconds of tone and light presentation, they were punished with a footshock produced by a shock stimulator (Med Associates, USA). After the offset of the tone, the rats could retrieve the reward without consequences. When rats retrieved the reward within 40 seconds without getting shocked (i.e. after tone offset), the trial was labelled as a successful trial. A shock trial was noted if the animal retrieved the reward during the 12 second tone presentation. If the rats did not retrieve the reward within 40 seconds in either NS or shock trials, the trial was labelled as an omission. After an omission, a new trial was started but a reward was only given when the food receptacle was empty. The rats were tested in daily RI sessions for three consecutive days followed by a rest day and then another six consecutive days. A shock intensity of 0.35 mA was used during the first three days. Next, the animals were tested for three days with a shock intensity of 0.45 mA, followed by three sessions with an intensity of 0.55 mA. These shock intensities were based on our previous study (Verharen et al., 2019), in which shock intensities were titrated for individual animals to reach a criterion of 20 success trials out of 30 stimulus trials, which required a median shock intensity of 0.50 mA (25-75^th^ percentile in between 0.40-0.60 mA).

#### Parameters

Successful reward collection, the number of omissions, shock count and reward collection latency in both stimulus and non-stimulus trials were used to evaluate overall performance in this task across the three different shock intensities.

### Elevated Plus Maze (EPM)

The EPM consisted of two open arms of 50 x 10 cm (L x W) and two closed arms of 50 x 10 x 40 cm (L x W x H) that extended from a central platform of 10 x 10 cm (L x W). The EPM was elevated to a height of 76 cm. It was located in a brightly lit room, with a light intensity of 300 lux on the open arms and central platform, and 240 lux on the closed arms. The test started with each rat being placed on the central platform facing an open arm. Testing lasted for 5 min and the maze was cleaned with water and soap between every trial. All trials were divided over two days, to prevent interference of short social isolation of the cage-mate. The uneven-numbered rats were tested on the first day and the even-numbered rats on the second day. The total time spent on the open arms, closed arms and central platform, the mean velocity and the total distance moved were assessed. All data was acquired through video recordings (Logitech C920 camera) and were analyzed with tracking software (EthoVision, Version 9.0.718, Noldus).

### Open Field (OF)

The OF was a circular arena with a diameter of 100 cm and 33.5 cm high walls. The OF was placed in a red-light-lit room. One external white light source was placed on the floor, which resulted in a light intensity of 12 lux in the OF. At the start of the trial, a rat was placed on a marked spot in the peripheral circle facing the walls. The test lasted for 10 min and the OF was cleaned with water and soap after each test. To avoid interference of social isolation, testing was divided over two days in the same way as mentioned for the EPM. Total time in the peripheral, middle and central zone, mean velocity, and total distances moved were measured. The peripheral, middle and central zones were created digitally in EthoVision (Version 9.0.718). Both the peripheral and central zones had a width of 20 cm, while the middle zones measured 10 cm. Data was obtained using video tracking software (EthoVision, Version 9.0.718, Noldus).

### Ex vivo electrophysiology

#### Slice preparation

Adult rats were anaesthetized after behavioural testing (P100 – P130) by induction with isoflurane and then transcardially perfused with ice-cold modified artificial cerebrospinal fluid (ACSF) containing (in mM): 92 Choline chloride, 2.5 KCl, 1.2 NaH_2_PO_4_, 30 NaHCO_3_, 20 HEPES, 25 glucose, 5 Na-ascorbate, 3 Na-pyruvate, 0.5 CaCl_2_.2H_2_O, and 10 MgSO_4_.7H_2_O, bubbled with 95% O_2_ and 5% CO_2_ (pH 7.3–7.4). The brain was quickly removed after decapitation and coronal slices of the mPFC (300 µm) were prepared using a vibratome (Leica VT1000S, Leica Microsystems) in ice-cold modified ACSF. Slices were initially incubated in the carbonated modified ACSF for 5 min at 35 °C and then transferred into a holding chamber containing standard ACSF containing (in mM): 126 NaCl, 3 KCl, 1.3 MgCl_2_.6H_2_O, 2 CaCl_2_.2H_2_O, 20 glucose, 1.25 NaH_2_PO_4_ and 26 NaHCO_3_ bubbled with 95% O_2_ and 5% CO_2_ (pH 7.3) at room temperature for at least 30 minutes. They were subsequently transferred to the recording chamber, perfused with standard ACSF that is continuously bubbled with 95% O_2_ and 5% CO_2_ at 28–32 °C.

#### Whole-cell recordings and analysis

Whole-cell patch-clamp recordings were performed from layer 5 pyramidal neurons in the mPFC. These neurons were visualized with an Olympus BX61W1 microscope using infrared video microscopy and differential interference contrast (DIC) optics. Patch electrodes were pulled from borosilicate glass capillaries and they had a resistance of 4-6 MΩ when filled with intracellular solutions. Action-potential independent miniature IPSCs (mIPSCs) were recorded in the presence of 6,7-dinitroquinoxaline-2,3-dione (DNQX) (20 µM) and D,L-2-amino-5-phosphopentanoic acid (D,L-AP5) (50 µM), with an internal solution containing (in mM): 70 K-gluconate, 70 KCl, 10 HEPES, 0.5 EGTA, 4 MgATP, 0.4 NaGTP, 4 Na2 phosphocreatine (pH 7.3 with KOH) in the presence of 1 μM tetrodotoxin (TTX) to block sodium channels. The membrane potential was held at -70 mV for voltage-clamp experiments. Signals were amplified, filtered at 2 kHz and digitized at 10 kHz using a MultiClamp 700B amplifier (Molecular Devices) and stored using pClamp 10 software. Series resistance was constantly monitored, and the cells were rejected from analysis if the resistance changed by >20% or it reached a value >30 MΩ. No series resistance compensation was used. Resting membrane potential was measured in bridge mode (I=0) immediately after obtaining whole-cell access. Passive and active membrane properties were analysed with Matlab (MathWorks) using a custom script and synaptic currents were analysed with Mini Analysis (Synaptosoft Inc., Decatur, GA). The detected currents were manually inspected to exclude false events.

### Data processing and statistical analyses

Statistical analyses were performed with GraphPad Prism (Software Inc.) and RStudio (R version 3.6.0. R Foundation for Statistical Computing). Data were analysed with a two-way ANOVA with housing condition and risky play as between-subjects factors followed by a post-hoc Tukey’s test where appropriate. In the electrophysiological data, the variance between cells within slices was larger than the variance between slices, indicating that individual cells can be treated as independent measurements. All graphs represent the mean ± standard error of the mean (SEM) with individual data points shown in coloured circles. The statistical range used in all figures: * p<0.05; ** p<0.01; *** p<0.001.

## Results

Rats were weaned on P21 and were subjected to either the control (CTL-LR), social play deprivation (SPD-LR), risky control (CTL-HR) or risky social play deprivation (SPD-HR) conditions, resulting in 4 groups of 12 rats. To assess the impact of high-risk social play on cognitive flexibility, a probabilistic reversal learning task (PRL) was used, which depends on PFC function (Fig. 2A,B (Dalton et al., 2016; Verharen et al., 2020; Bijlsma et al., 2022). Male rats in the high-risk groups (CTL-HR and SPD-HR) reached a higher performance level in terms of rewards obtained compared to the low-risk groups (CTL-LR and SPD-LR) (Fig. 2C), mainly caused by a difference between the CTL-LR and CTL-HR animals. All male groups completed a comparable amount of reversals (Fig. 2D). The difference in performance could be explained by an enhanced tendency of the high-risk animals to stay at a lever after it was rewarded (win-stay behaviour; Fig. 2E). Consistent with our previous findings (Bijlsma et al., 2022), male social play-deprived rats displayed enhanced win-stay behaviour compared to their socially housed counterparts, but this was only the case within the low-risk groups. In female rats, the high-risk groups obtained more rewards (Fig. 2F), achieved more reversals (Fig. 2G) and showed a higher percentage of win-stay behaviour (Fig. 2H) in comparison to their low-risk counterparts. Additionally, social play-deprived females (SPD-LR and SPD-HR) obtained more rewards than control females (CLT-LR and CTL-HR) (Fig. 2F) while achieving a comparable amount of reversals and showing similar win-stay behaviour (Fig. 2G,H). Please note that performance of females and males cannot be directly compared, as females received less trials in accordance with their smaller body weight (see methods). Our observations indicate that cognitive performance in adult rats was altered after the opportunity to take risks during social play in both male and female rats. SPD affected win-stay behaviour of males and the number of rewards collected in females, which is in general agreement with our previous findings (Bijlsma et al., 2022).

**Figure 2.**
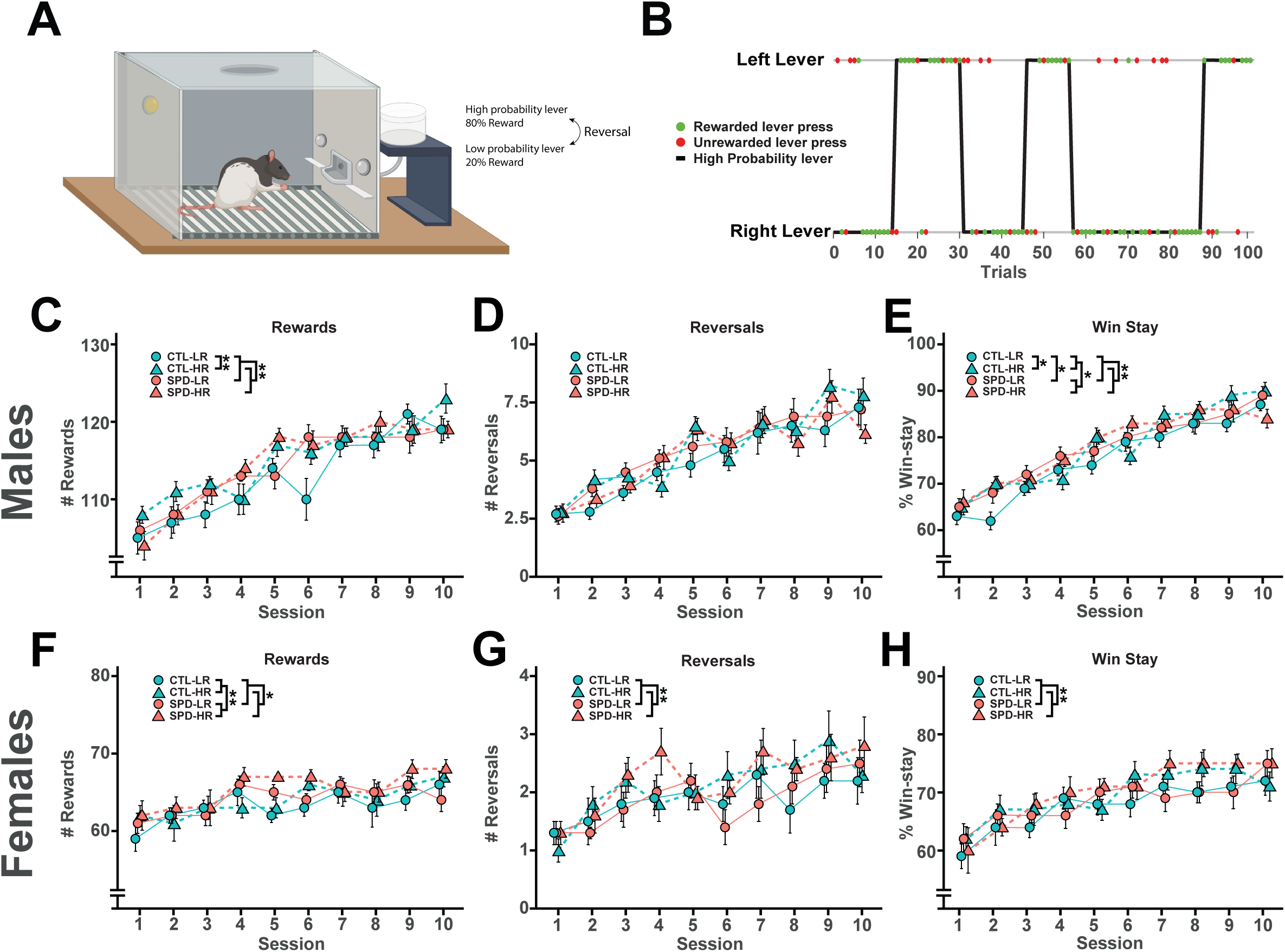
Performance in the PRL task after social play deprivation with or without the opportunity for risky play. (A) PRL task. (B) Representation of the 100 lever presses of an example session. Green and red dots represent rewarded and unrewarded lever presses respectively. A reversal is indicated by the change of the high probability lever. (C-E) PRL results from male rats. (C) Rewards obtained per session (2W-ANOVA, Housing: p=0.26, Risk: p=0.003, Interaction: p=0.024; TUKEY, CTLLR-SPDLR: p=0.18, CTLLR-CTLHR: p=0.005, SPDLR-SPDHR: p=0.84, SPDHR-CTLHR: p=0.95). (D) Reversals achieved per session (2W-ANOVA, Housing: p=0.41, Risk: p=0.35, Interaction: p=0.083). (E) % Win-stay behaviour per session (2W-ANOVA, Housing: p=0.035, Risk: p=0.010, Interaction: p=0.039; TUKEY, CTLLR-SPDLR: p=0.048, CTLLR-CTLHR: p=0.022, SPDLR-SPDHR: p=0.86, SPDHR-CTLHR: p=0.99). (F-H) PRL results from female rats. (F) Rewards obtained per session (2W-ANOVA, Housing: p=0.003, Risk: p=0.049, Interaction: p=0.89). (G) Reversals achieved per session (2W-ANOVA, Housing: p=0.68. Risk: p=0.006, Interaction: p=0.81). (H) % Win-stay behaviour per session (Circles, 2W-ANOVA, Housing: p=0.23, Risk: p=0.006, Interaction: p=0.99). Statistical range (p): * p<0.05; ** p<0.01; *** p<0.001.

Behavioural control was assessed using a response inhibition (RI) task, in which rats need to wait to collect a reward in half of the trials (indicated by a light and tone cue), with foot shocks of increasing intensity as a penalty (Fig. 3A) (Verharen et al., 2019). During stimulus trials, male rats in all groups showed a higher success rate (thus avoiding punishment and consequently receiving less shocks) when the foot shock intensity was increased (Fig. 3B-C). Rats in the high-risk groups succeeded more often to collect their rewards (Fig. 3B), and received fewer shocks than low-risk rats (Fig. 3C). The number of omissions increased with higher shock intensity and was comparable between the groups (Fig. 3D). Deprivation of social play behaviour did not affect RI task performance in male rats. SPD affected the latency of reward retrieval only during non-stimulus trials with SPD animals collecting the reward faster than CTL rats (Table 1). No differences in the number of rewards retrieved or omissions during non-stimulus trials were found between groups (data not shown). Females tended to show more avoidance behaviour in the RI task by not approaching the reward dispenser (and thus made more omissions)(Svoboda et al., 2012; Yokota et al., 2017; Finnell et al., 2018). In female rats, there were no effects of SPD or the opportunity for risky play on performance in the RI task. Similar to the males, the success rate for female rats increased when the shock intensity was increased (Fig. 3E) and the number of shocks received decreased (Fig. 3F). The number of omissions did not change with shock intensity (Fig. 3G). No effect was found on reward retrieval latency in both male and female rats with an exception that female high-risk rats had a lowered reward retrieval during non-stimulus trials (data not shown; (2W-ANOVA, Housing: p=0.38, Risk: p=0.004, Interaction: p=0.17). No differences in the number of rewards retrieved or omissions during non-stimulus trials were found between groups (data not shown). Our data suggest that male, but not female rats, that engaged in risky play as juveniles (i.e., the CTL-HR and SPD-HR groups) showed better behavioural inhibition in this task, whereas deprivation of social play did not affect RI task performance. (Svoboda et al., 2012; Avcu et al., 2014; Shanazz et al., 2022),

**Figure 3.**
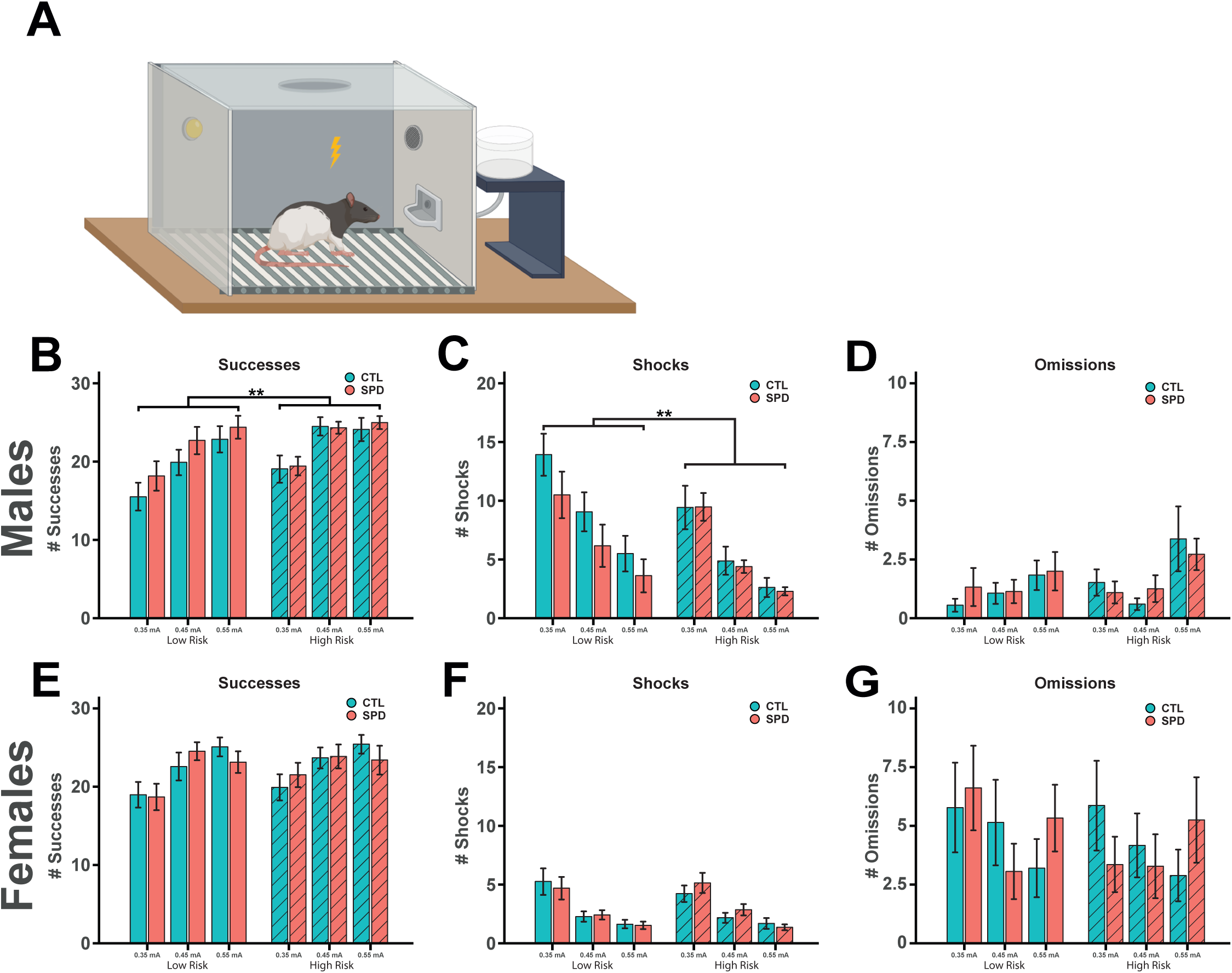
Performance in the RI task after social play deprivation with or without the opportunity for risky play. RI task (A) results for male (B-D) and female rats (E-G). (B) Successes during stimulus trials (2W-ANOVA, Housing: p=0.084, Risk: p=0.004, Interaction: p=0.18, Shock Intensity: p<0.001; TUKEY, 0.35mA – 0.45mA: p<0.001, 0.35mA – 0.55mA: p<0.001, 0.45mA – 0.55mA: p=0.45). (C) The number of shocks received during stimulus trials (2W-ANOVA, Housing: p=0.076, Risk: p=0.001, Interaction: p=0.12, Shock Intensity: p<0.001; TUKEY, 0.35mA – 0.45mA: p<0.001, 0.35mA – 0.55mA: p<0.001, 0.45mA – 0.55mA: p=0.021). (D) The number of omissions stimulus trials (2W-ANOVA, Housing: p=0.66, Risk: p=0.072, Interaction: p=0.34, Shock Intensity: p<0.001; TUKEY, 0.35mA – 0.45mA: p=0.89, 0.35mA – 0.55mA: p<0.001, 0.45mA – 0.55mA: p<0,001). (E) Successes during stimulus trials (2W-ANOVA, Housing: p=0.88, Risk: p=0.24, Interaction: p=0.98, Shock Intensity: p<0.001; TUKEY, 0.35mA – 0.45mA: p<0.001, 0.35mA – 0.55mA: p<0.001, 0.45mA – 0.55mA: p=0.72). (F) The number of shocks received during the 30 stimulus trials (2W-ANOVA, Housing: p=0.68, Risk: p=0.84, Interaction: p=0.33, Shock Intensity: p<0.001; TUKEY, 0.35mA – 0.45mA: p<0.001, 0.35mA – 0.55mA: p<0.001, 0.45mA – 0.55mA: p=0.053). (G) The number of omissions during stimulus trials (2W-ANOVA, Housing: p=0.97, Risk: p=0.29, Interaction: p=0.64, Shock Intensity: p=0.17). Statistical range (p): * p<0.05; ** p<0.01; *** p<0.001.

**Table 1:** Schematic overview of main effects of social play deprivation and the opportunity for risky play on male and female performance and behaviour.

|  |  | *Risky play* |  | *Social play deprivation* |  |
| --- | --- | --- | --- | --- | --- |
|  |  | Males | Females | Males | Females |
| *PRL* | Rewards | ↑ | ↑ |  | ↑ |
|  | Reversals |  | ↑ |  |  |
|  | Win-Stay | ↑ | ↑ | ↑ |  |
| *RI* | Successes | ↑ |  |  |  |
|  | Shocks | ↓ |  |  |  |
|  | Omissions |  |  |  |  |
| *EPM* | Open | ↓ |  |  | ↑ |
|  | Closed |  |  |  | ↓ |
|  | Center |  |  |  |  |
|  | Distance |  |  |  |  |
| *OF* | Center |  | ↑ |  |  |
|  | Middle |  |  |  |  |
|  | Periphery |  |  |  |  |
| *IPSCs* | Frequency | ↑ |  | ↓ | ↓ |
|  | Amplitude | ↑ |  |  |  |
|  | Rise-time | ↓ | ↓ |  |  |
|  | Decay-time |  |  |  |  |

“↑” increased/improved/higher, “↓” decreased/impaired/lower compared to control animals

To assess anxiety-like behaviour and locomotor activity of the rats, we used the elevated plus maze (EPM, Fig. 4A) and the open field (OF). Male rats in the high-risk groups spent less time on the open arms (Fig. 4B) and showed a tendency towards more time on the closed arms of the EPM (Fig. 4C) compared to the low-risk animals. Time spent in the centre of the EPM did not differ between the groups (Fig. 4D). No differences were found between CTL and SPD groups, and locomotion on the EPM was comparable between the four groups of male rats (Fig. 4E). We did not find differences between the low- and high-risk groups in the female rats, but social play-deprived females spent more time on the open arms (Fig. 4F) and less time on the closed arms of the EPM (Fig. 4G) compared to control rats. No differences between groups were found in the time spent in the centre of the maze (Fig. 4H). All female groups showed comparable locomotor activity on the EPM (Fig. 4I).

**Figure 4.**
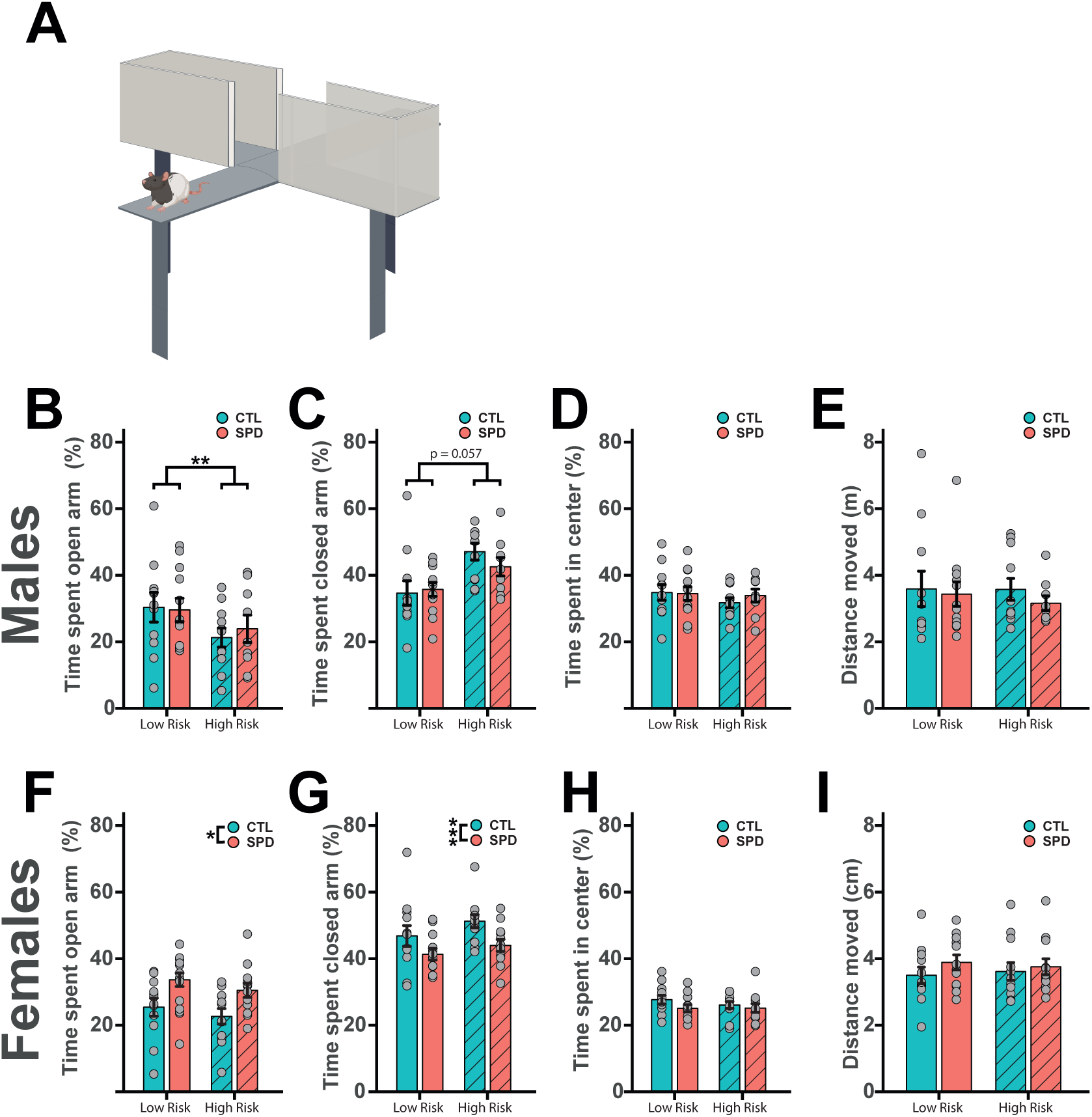
Behaviour on the EPM after social play deprivation with or without the opportunity for risky play. EPM results (A) for male (B-E) and female (F-I) rats. (B) Total distance moved (2W-ANOVA, Housing: p=0.50, Risk: p=0.73, Interaction: p=0.75). (C-E) Percentage time spent on the (C) open arms (2W-ANOVA, Housing: p=0.39, Risk: p=0.002, Interaction: p=0.30), (D) closed arms (2W-ANOVA, Housing: p=0.71, Risk: p=0.057, Interaction: p=0.74) and (E) in the centre zone (2W-ANOVA, Housing: p=0.68, Risk: p=0.35, Interaction: p=0.67). (F) Total distance moved (2W-ANOVA, Housing: p=0.29, Risk: p=0.97, Interaction: p=0.62). (G-I) Percentage time spent on the (G) open arms (2W-ANOVA, Housing: p=0.019, Risk: p=0.089, Interaction: p=0.45), (H) closed arms (2W-ANOVA, Housing: p<0.001, Risk: p=0.27, Interaction: p=0.78) and (I) in the centre zone (2W-ANOVA, Housing: p=0.27, Risk: p=0.81, Interaction: p=0.79). Statistical range (p): * p<0.05; ** p<0.01; *** p<0.001.

Behaviour of the male rats in the OF (Fig. 5A) was unaffected by play deprivation or the opportunity to take risks during play. Similar to the EPM, no differences were found in the distance travelled (Fig. 5B). Additionally, time spent in the centre, middle and periphery was comparable between all groups (Fig. 5C-E). Female rats are generally more active than male rats (Svoboda et al., 2012; Avcu et al., 2014; Shanazz et al., 2022), presumably because they are smaller. Consistently, we observed that females showed higher locomotion in the OF (Scholl et al., 2019). The behaviour of the female rats in the OF was also not affected by play deprivation or the opportunity to take risks during play (Fig. 5F-I) with the exception that CTL-HR rats spent more time in the centre (Fig. 5G) and less time in the periphery (Fig. 5I) in comparison to their CTL-LR counterparts. In summary, we observed no effect on the locomotor activity of the rats by SPD or risk-taking during play in these tests. Anxiety-like behaviour, however, was affected by risk-taking in male, but not female, rats while SPD affected anxiety-like behaviour in female, but not male, rats.

**Figure 5.**
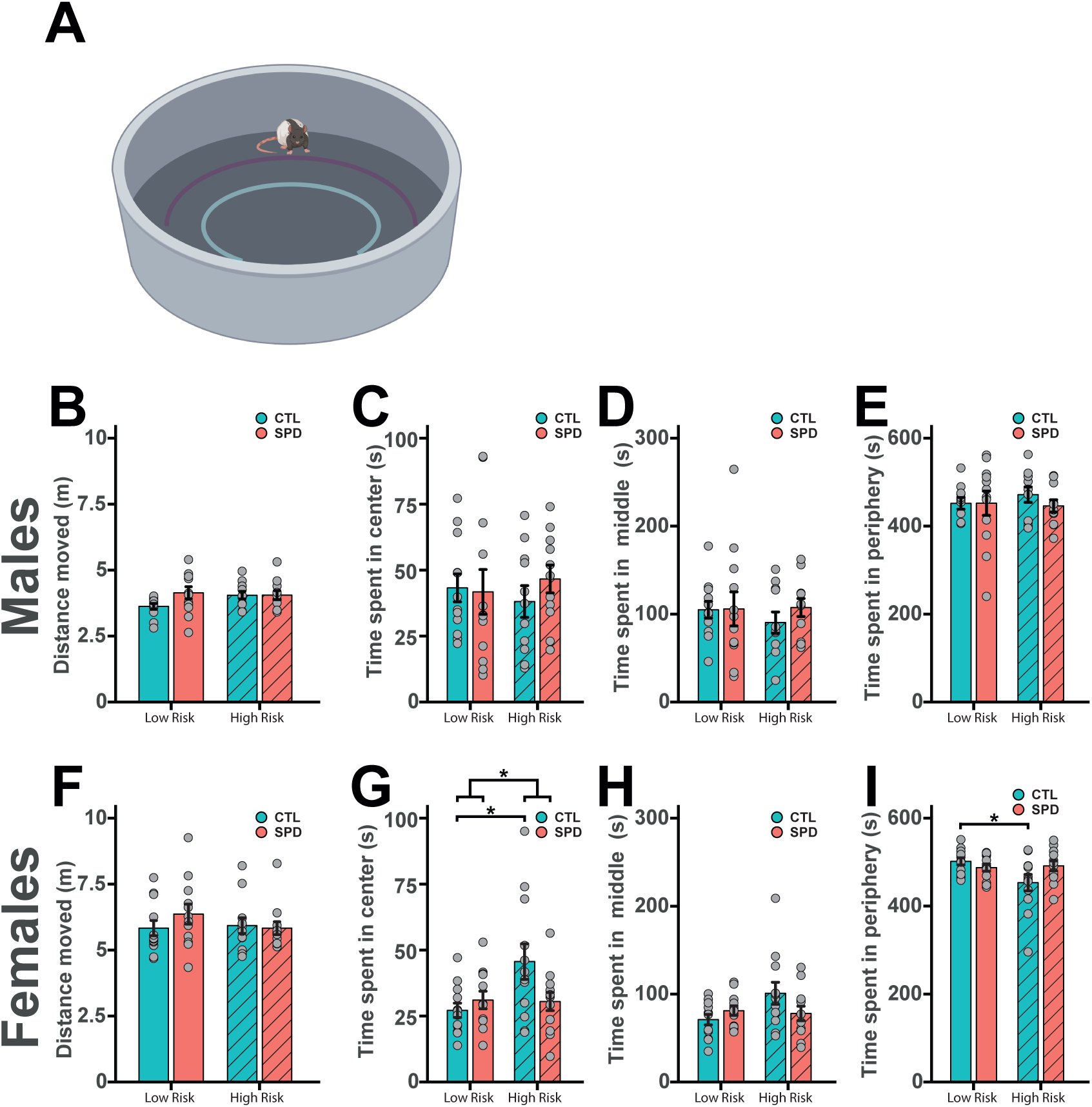
Behaviour in the OF after social play deprivation with or without the opportunity for risky play. OF results (A) for male (B-E) and female (F-I) rats. (B) Total distance moved (2W-ANOVA, Housing: p=0.12, Risk: p=0.33, Interaction: p=0.14). (C-E) Time spent in the (C) centre (2W-ANOVA, Housing: p=0.61, Risk: p=0.98, Interaction: p=0.44), (D) middle (2W-ANOVA, Housing: p=0.53, Risk: p=0.64, Interaction: p=0.55) and (E) periphery (2W-ANOVA, Housing: p=0.53, Risk: p=0.72, Interaction: p=0.50). (F) Total distance moved (2W-ANOVA, Housing: p=0.48, Risk: p=0.48, Interaction: p=0.31). (G-I) Time spent in the (G) centre (2W-ANOVA, Housing: p=0.20, Risk: p=0.046, Interaction: p=0.034; TUKEY, CTLLR-SPDLR: p=0.92, CTLLR-CTLHR: p=0.022, SPDLR-SPDHR: p=0.99, SPDHR-CTLHR: p=0.081), (H) middle (2W-ANOVA, Housing: p=0.46, Risk: p=0.12, Interaction: p=0.058) and (I) periphery (2W-ANOVA, Housing: p=0.34, Risk: p=0.078, Interaction: p=0.041; TUKEY, CTLLR-SPDLR: p=0.85, CTLLR-CTLHR: p=0.039, SPDLR-SPDHR: p=0.99, SPDHR-CTLHR: p=0.14). Statistical range (p): * p<0.05; ** p<0.01; *** p<0.001.

To study the impact of the opportunity to take risks during play and social deprivation on the development of PFC inhibitory circuitry, we performed voltage-clamp recordings from layer 5 pyramidal cells of the mPFC (Fig. 6A,B) of male (Fig. 6C-F) and female (Fig. 6G-J) rats. Consistent with our previous findings (Bijlsma et al., 2022, 2023), a reduction in the frequency of miniature inhibitory postsynaptic currents (mIPSCs) was found in male SPD in comparison with CTL rats (Fig. 6C). We observed an increase in mIPSC frequency in the male high-risk groups in comparison with their low-risk counterparts (Fig. 6C). The mIPSCs in the high-risk groups also had higher amplitudes (Fig. 6D) and faster rise times (Fig. 6E). Decay kinetics (Fig. 6F) were comparable between all groups. In contrast to the males, we found no difference in mIPSC frequency between the low and high-risk groups in female rats (Fig. 6G). However, the reduction in mIPSC frequency in SPD compared to CTL was consistent with our previous observations in males (Fig. 6G). Amplitudes of the measured events were comparable between all groups (Fig. 6H). Similar to our observation in male rats, mIPSC rise time kinetics were faster in the high-risk group (Fig. 6I) while decay kinetics remained unaffected (Fig. 6J). These results confirm that SPD reduces the frequency of mIPSCs in layer 5 pyramidal cells of the mPFC (Bijlsma et al., 2022, 2023), and extend this observation to female rats. The opportunity to take risks increased mIPSC frequency in males, but not females, while faster mIPSC rise times in the high-risk groups were observed in both sexes.

**Figure 6.**
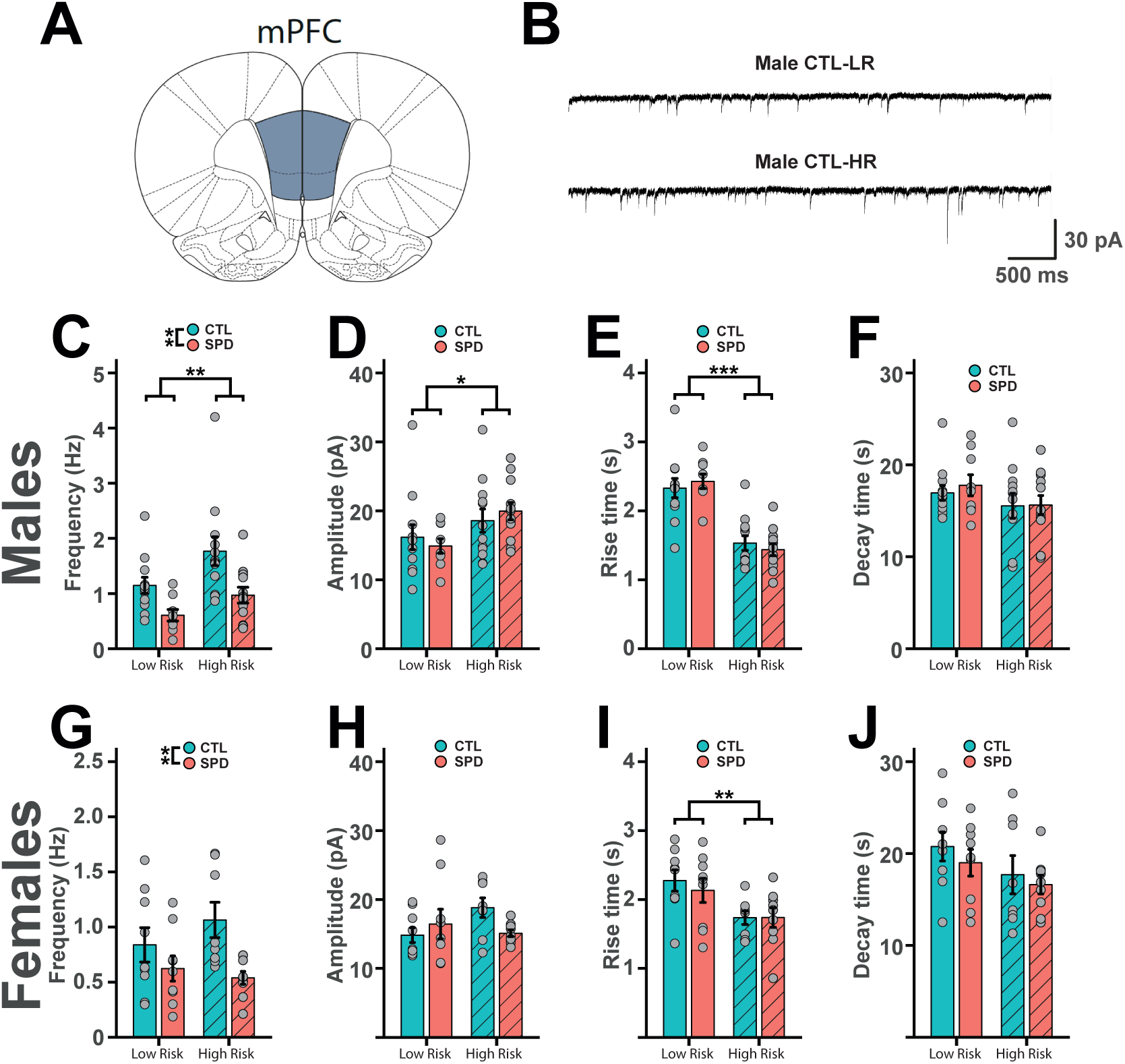
Prefrontal inhibition in L5 of the mPFC after social play deprivation with or without the opportunity for risky play. (A) Schematic diagram depicting the recording site in the mPFC. (B) Example traces of miniature IPSCs (mIPSCs) in L5 pyramidal cells in slices from control CTL-LR and CTL-HR male rats. Results for male (C-F) and female (G-J) rats. (C) Frequency (2W-ANOVA, Housing: p=0.001, Risk: p=0.009, Interaction: p=0.49), (D) amplitude (2W-ANOVA, Housing: p=0.79, Risk: p=0.024, Interaction: p=0.39), (E) rise time (2W-ANOVA, Housing: p=0.54, Risk: p<0.001, Interaction: p=0.40) and (F) decay time (2W-ANOVA, Housing: p=0.79, Risk: p=0.12, Interaction: p=0.74) of mIPSC events. (G) Frequency (2W-ANOVA, Housing: p=0.007, Risk: p=0.61, Interaction: p=0.23), (H) amplitude (2W-ANOVA, Housing: p=0.51, Risk: p=0.39, Interaction: p=0.069), (I) rise time (2W-ANOVA, Housing: p=0.56, Risk: p=0.004, Interaction: p=0.63) and (J) decay time (2W-ANOVA, Housing: p=0.35, Risk: p=0.089, Interaction: p=0.83) of mIPSC events. Statistical range (p): * p<0.05; ** p<0.01; *** p<0.001.

We summarize all results in Table 1. Together, our data indicate that the opportunity to take risks during play alters cognitive flexibility in both males and females and improved behavioural control in a response inhibition task in male rats. Anxiety-like behaviour was only modestly altered by risk. Deprivation of social play behaviour altered cognitive flexibility in both sexes but had no consequences for response inhibition. Anxiety-like behaviour was modestly reduced only in female rats after SPD. Our findings also demonstrate that the reduction in inhibitory synaptic inputs in the mPFC of rats by SPD is present in both male and female rats while the opportunity to play in a high-risk environment increased mIPSC frequency only in male rats. In both sexes, mIPSCs displayed faster rise times in high risk animals.

## Discussion

In this study, we assessed how two important aspects of play, e.g. social play with peers and the opportunity for risk-taking during play, affect cognitive and emotional behaviour and PFC function in adulthood. Our results suggest that these two aspects of play both affect PFC circuitry and adult behaviour, but with limited overlap. We show that providing animals with the possibility to play in a high-risk environment during a peak social play period (P21-P42) altered cognitive flexibility and anxiety-like behaviour in both sexes. Behavioural inhibition was improved only in males. At the neuronal level, we found that the frequency of mIPSCs was increased in male rats that engaged in play in a high-risk environment. The mIPSC frequency increase was not present in female rats, but mIPSCs had faster rise kinetics in high-risk animals in both sexes. SPD altered cognitive flexibility and reduced mIPSC frequency in males and females, which is in general agreement with our previous findings (Bijlsma et al., 2022, 2023). In addition, we found that response inhibition was unaffected by SPD, while anxiety-like behaviour was affected only in female rats.

As stated in the Introduction, in this study we aimed to answer four questions. The first of these was how the opportunity for risk-taking behaviour during play alters the development of cognitive flexibility, behavioural inhibition and anxiety-like behaviour. To the best of our knowledge, our study is the first to explicitly address risky play in an animal model. Of note, our approach bears some resemblance to previous studies using environmental enrichment, the main difference being that our ‘enrichment’ was tailored to emulate aspects of risky play as described by Sandseter (2007). In addition, unlike most enrichment experiments, in which animals are housed in this environment throughout the study, we restricted access to the risky play environment to two 30-min periods per day during the three weeks in life when play is most abundant in rats (i.e. P21-42; Panksepp, 1981), and performed behavioural testing several weeks later. Thus, our approach focused on the long-term developmental consequences of high-risk play during post-weaning development, rather than its acute effects on behaviour and PFC function. We found that the opportunity for risk-taking behaviour led to altered cognitive flexibility in both male and female rats in the PRL task, shown by the higher percentage of win-stay behaviour and rewards earned during a session. Furthermore, in high-risk female rats, an increase in the number of reversals achieved was also found. Comparable changes in cognitive flexibility have been described in similar reversal learning designs after enriched or complex housing (Sampedro-Piquero et al., 2015; Zeleznikow-Johnston et al., 2017). At face value, this pattern of changes indicates that exposure to a high-risk play environment enhances later cognitive flexibility, as increases in the number of rewards and reversals are typically interpreted as an improvement in task performance(see e.g. Dalton et al., 2016; Verharen et al., 2020).However, in a previous study (Bijlsma et al., 2022) we found that in this PRL setup achieving more reversals, rewards and increased win-stay behaviour can also be the result of a simplified cognitive strategy. Therefore, we need to be cautious with a straightforward ‘better-or-worse’ performance interpretation here.

When we assessed response inhibition, we observed that male, but not female, rats that had opportunities for risk-taking during play were better able to wait during stimulus trials. As a result, they achieved more successes and received fewer shocks than their low-risk counterparts. Performance in this behavioural inhibition task consists of multiple components like control over behaviour, appreciation of stimulus value and task engagement (Verharen et al., 2019). Our results indicate an increase in control over behaviour after risk-taking as the high-risk male rats were better in refraining from taking the sucrose pellet during stimulus presentation. The stimulus value seems not to be affected as there were clear differences between reward collection latency during stimulus and non-stimulus trials, with no effect of risky play. Furthermore, as omissions were not affected, we think that task engagement between all groups was comparable. In sum, engaging in risky play when young clearly affects cognitive development, as altered cognitive flexibility was seen in both male and female rats, and enhanced behavioural inhibition in males.

In this study we also show that anxiety-like behaviour is affected by risk-taking in male rats as seen in by a reduction of time spent on the open arms of the EPM. Multiple studies have examined the effects of enrichment of the housing condition on anxiety related behaviour in rats, but mixed effects on behaviour in the EPM have been reported. That is, after enrichment rats have been reported to spend more time on the open arms of the EPM (Friske and Gammie, 2005; Baldini et al., 2013), less time on the open arms (Branchi and Alleva, 2006; Sparling et al., 2018) or no effect of enrichment was found at all (Brenes et al., 2009; Green et al., 2010; Li et al., 2016). As mentioned above, an important difference with our study is that in most, if not all of these studies, behavioural testing was done when the animals were still housed in their enriched environment, whereas in our study the rats had the opportunity to play in an enriched environment only when they were juveniles. In addition, our high-risk rats are well familiar with moving, climbing and playing on elevated platforms as our risky-play cage allowed them to do exactly this. The reduction in time spent on the open arms in male rats after high-risk play that we observed may therefore reflect improved risk assessment. Interestingly, the effect of risky play was only noticeable in females in the OF, since female CTL-HR rats spent more time in the centre of the OF. In short, the opportunity to take risks during play when young affects anxiety-like behaviour in both males and females, but in a somewhat different fashion.

The second question was how social play experience affects the development of cognitive flexibility, behavioural inhibition and anxiety-like behaviour. We found that SPD affected the win-stay behaviour of males and the number of rewards collected in females in the PRL task, comparable to the effects of SPD on PRL performance in our previous work (Bijlsma et al., 2022). These results also resonate well with previous studies that showed that alterations in the social domain during the post-weaning period resulted in impairments in impulsive actions and decision-making under challenging and novel circumstances (Baarendse et al., 2013; Schneider et al., 2016). As we did not observe effects of SPD on response inhibition, the effects of SPD on executive functions appears to be specific to the cognitive domain of flexibility. While no effects of SPD were found on the behaviour on the EPM in male rats, socially deprived female rats spent more time in the open arms and less time in the closed arms. This was not expected as previous studies have shown that play deprivation in rats can increase anxiety-related responses (Wright et al., 1991; Arakawa, 2005; Lukkes et al., 2009). However, a lack of effect of SPD on anxiety-related behaviour has also been reported (Van den Berg et al., 1999), which is consistent with our null findings in the OF. Although speculative, the apparent reduction in anxiety-like behaviour on the EPM in female SPD rats may reflect difficulty in coping with a challenging environment, reflecting increased risk taking.

The third question addressed how play manipulation affects the maturation of the mPFC. We previously showed that SPD results in a specific reduction of inhibitory currents in L5 cells in the adult mPFC, while excitatory currents remained unaffected (Bijlsma et al., 2022, 2023). Here we observed this reduction in mIPSC frequency after SPD in both the low-risk and high-risk groups and it was also present in female rats. This emphasizes the general importance of post-weaning social play, consistent with earlier studies that identified the time window between weaning and puberty (P21-P42) as a critical period for PFC maturation (Lukkes et al., 2009; Kolb et al., 2012; Baarendse et al., 2013; Himmler et al., 2018; Bijlsma et al., 2022, 2023). In addition, we found that the frequency of mIPSCs in L5 cells was increased in the high-risk groups in comparison to the low-risk groups, but only in male rats. In slices from both sexes, mIPSCs showed faster rise kinetics in the high-risk group. The combination of increased mIPSC frequency and enhanced kinetics suggest an increase in perisomatic synapses after high-risk play. We previously showed that SPD affects perisomatic synapses made by parvalbumin (PV)-positive interneurons (Bijlsma et al., 2022). We cannot exclude (partial) overlap between the synapses affected, but our data show that the synaptic alterations after high-risk play and social play experience (i.e. that are counteracted by SPD) are largely additive. These findings therefore suggest that SPD and the opportunity for risk-taking have independent effects on the development of IPSCs in the PFC.

Finally, our data addresses the question if the consequences of risky play and SPD differ between males and females. In the PRL task, exposure to a high-risk play environment and SPD had largely comparable effects in male and female rats. A higher rate of avoidance behaviour and a lower shock threshold for females have been reported before (Svoboda et al., 2012; Yokota et al., 2017; Finnell et al., 2018), which in the our RI task, causes female animals to use a different strategy compared to males, but not at the cost of earning less rewards. Exposure to a high-risk play environment improved RI task performance in male rats only, suggesting that high-risk play differentially influenced the different strategies used by male and female rats in this task. Similarly, the effect of high-risk play on the behaviour in the EPM was solely found in males, although in this case, high-risk male animals showed more (rather than less) anxiety-like behaviour, perhaps reflecting increased risk assessment. When comparing the behaviour of the two sexes on the EPM, females tended to spend more time in the closed arms and less in the centre zone than males, consistent with a higher level of avoidance behaviour. This trend was also seen in the OF, where females spent less time in the centre and middle areas of the open field, preferring to stay more in the periphery of the arena. SPD resulted in less anxiety-like behaviour only in female rats, perhaps as a result of more risky-taking or a lower level of avoidance behaviour, whereas in the OF, high-risk play reduced anxiety-like behaviour in females only. Our results indicate that the effects of SPD and high-risk play on task performance are both sex-dependent and strategy-dependent. The consequences at the neurobiological level appear mostly independent of sex, as at the decrease in mIPSC frequency after SPD and faster risetimes after high-risk play were observed in both sexes. However, high-risk play resulted in an increase in mIPSC frequency that was specific for males. Future studies will need to clarify if specific connections are affected. Multiple enrichment studies have previously described sex differences in behavioural tasks after exposure to a challenging environment (Elliott and Grunberg, 2005; Simpson and Kelly, 2011; Chamizo et al., 2016). However, we are the first to describe sex differences after the opportunity to take risks during social play in combination with a play deprivation paradigm.

Together, our data show that exposure to risks during juvenile play, and the opportunity to play with peers affect performance in PFC-dependent cognitive tasks and anxiety-like behaviour in adulthood, as well as the development of inhibition in the mPFC. Some of the consequences of risky and social play were different between the sexes. Furthermore, our results indicate that interactions between risky play and SPD were limited, suggesting that social play with peers and the opportunity for risk-taking during play independently affect the development of behaviour and PFC function. This implies that the effects of SPD cannot be mitigated by increasing the opportunity for risk-taking, but that both aspects of juvenile play have separate value for adult behaviour.

## Author Contribution

E.J. Marijke Achterberg (MA), Annemarie J.M. Baars (AMB), Ate Bijlsma (AB), Evelien E. Birza (EB), Marieke J.J.M van Gaans (MG), Heidi M.B. Lesscher (HL), José G. Lozeman-van t Klooster (JK), Janneke Maranus (JM), Tara C. Pimentel (TP), Louk J.M.J. Vanderschuren (LV), Corette J. Wierenga (CW)

The study was conceived by LV, CW, HL, and MA. Experiments were designed by AB, EB, TP, JK, AMB, CW, LV, HL, and MA. Experiments were performed by AB, EB, TP, JM, and MG. Analysis was performed by AB and EB. The manuscript was written by AB, CW, LV, HL, and MA with input from all other authors.

## References

Allen and Rapee (2005) Anxiety Disorders. In: Cognitive Behaviour Therapy for Children and Families, 2nd ed. (P. Graham, ed), pp 300–319. Cambridge University Press.

Arakawa H (2005) Interaction between isolation rearing and social development on exploratory behavior in male rats. Behav Processes 70:223–234.

Avcu P, Jiao X, Myers CE, Beck KD, Pang KCH, Servatius RJ (2014) Avoidance as expectancy in rats: Sex and strain differences in acquisition. Front Behav Neurosci 8:1–8.

Baarendse PJJ, Counotte DS, O’Donnell P, Vanderschuren LJMJ (2013) Early social experience is critical for the development of cognitive control and dopamine modulation of prefrontal cortex function. Neuropsychopharmacology 38:1485–1494.

Baarendse PJJ, Limpens JHW, Vanderschuren LJMJ (2014) Disrupted social development enhances the motivation for cocaine in rats. Psychopharmacology (Berl) 231:1695–1704.

Baldini S, Restani L, Baroncelli L, Coltelli M, Franco R, Cenni MC, Maffei L, Berardi N (2013) Enriched early life experiences reduce adult anxiety-like behavior in rats: A role for insulin-like growth factor 1. J Neurosci 33:11715–11723.

Bari A, Theobald DE, Caprioli D, Mar AC, Aidoo-Micah A, Dalley JW, Robbins TW (2010) Serotonin modulates sensitivity to reward and negative feedback in a probabilistic reversal learning task in rats. Neuropsychopharmacology 35:1290–1301 Available at: 10.1038/npp.2009.233.

Bell HC, McCaffrey DR, Forgie ML, Kolb B, Pellis SM (2009) The Role of the Medial Prefrontal Cortex in the Play Fighting of Rats. Behav Neurosci 123:1158–1168.

Bijlsma A, Omrani A, Spoelder M, Verharen JPH, Bauer L, Cornelis C, de Zwart B, van Dorland R, Vanderschuren LJMJ, Wierenga CJ (2022) Social play behavior is critical for the development of prefrontal inhibitory synapses and cognitive flexibility in rats. J Neurosci 00:JN-RM-0524-22.

Bijlsma A, Vanderschuren LJMJ, Wierenga CJ (2023) Social play behavior shapes the development of prefrontal inhibition in a region-specific manner. Cereb Cortex:1–10.

Branchi I, Alleva E (2006) Communal nesting, an early social enrichment, increases the adult anxiety-like response and shapes the role of social context in modulating the emotional behavior. Behav Brain Res 172:299–306.

Brenes JC, Padilla M, Fornaguera J (2009) A detailed analysis of open-field habituation and behavioral and neurochemical antidepressant-like effects in postweaning enriched rats. Behav Brain Res 197:125–137.

Brussoni M, Olsen LL, Pike I, Sleet DA (2012) Risky play and children’s safety: Balancing priorities for optimal child development. Int J Environ Res Public Health 9:3134–3148.

Chamizo VD, Rodríguez CA, Sánchez J, Mármol F (2016) Sex differences after environmental enrichment and physical exercise in rats when solving a navigation task. Learn Behav 44:227–238 Available at: 10.3758/s13420-015-0200-3.

Dalton GL, Wang NY, Phillips AG, Floresco SB (2016) Multifaceted contributions by different regions of the orbitofrontal and medial prefrontal cortex to probabilistic reversal learning. J Neurosci 36:1996–2006.

Elliott BM, Grunberg NE (2005) Effects of social and physical enrichment on open field activity differ in male and female Sprague-Dawley rats. Behav Brain Res 165:187–196.

Finnell JE, Muniz BL, Padi AR, Lombard CM, Moffitt CM, Wood CS, Wilson LB, Reagan LP, Wilson MA, Wood SK (2018) Essential Role of Ovarian Hormones in Susceptibility to the Consequences of Witnessing Social Defeat in Female Rats. Biol Psychiatry 84:372–382.

Friske JE, Gammie SC (2005) Environmental enrichment alters plus maze, but not maternal defense performance in mice. Physiol Behav 85:187–194.

Graham KL, Burghardt GM (2010) Current perspectives on the biological study of play: Signs of progress. Q Rev Biol 85:393–418.

Gray P (2011) The Decline of Play and the Rise of Psychopathology in Children and Adolescents. Am J Play 3:443–463 Available at: http://www.psychologytoday.com/files/attachments/1195/ajp-decline-play-published.pdf.

Gray P (2017) What exactly is play, and why is it such a powerful vehicle for learning? Top Lang Disord 37:217– 228.

Green TA, Alibhai IN, Roybal CN, Winstanley CA, Theobald DEH, Birnbaum SG, Graham AR, Unterberg S, Graham DL, Vialou V, Bass CE, Terwilliger EF, Bardo MT, Nestler EJ (2010) Environmental Enrichment Produces a Behavioral Phenotype Mediated by Low Cyclic Adenosine Monophosphate Response Element Binding (CREB) Activity in the Nucleus Accumbens. Biol Psychiatry 67:28–35 Available at: 10.1016/j.biopsych.2009.06.022.

Himmler BT, Mychasiuk R, Nakahashi A, Himmler SM, Pellis SM, Kolb B (2018) Juvenile social experience and differential age-related changes in the dendritic morphologies of subareas of the prefrontal cortex in rats. Synapse 72:1–9.

Kolb B, Mychasiuk R, Muhammad A, Li Y, Frost DO, Gibb R (2012) Experience and the developing prefrontal cortex. Proc Natl Acad Sci U S A 109:17186–17196.

Lavrysen A, Bertrands E, Leyssen L, Smets L, Vanderspikken A, De Graef P (2017) Risky-play at school. Facilitating risk perception and competence in young children. Eur Early Child Educ Res J 25:89–105.

Lesscher HMB, Spoelder M, Rotte MD, Janssen MJ, Hesseling P, Lozeman-Van’t Klooster JG, Baars AM, Vanderschuren LJMJ (2015) Early social isolation augments alcohol consumption in rats. Behav Pharmacol 26:673–680.

Li KA, Lund ET, Voigt JPW (2016) The impact of early postnatal environmental enrichment on maternal care and offspring behaviour following weaning. Behav Processes 122:51–58.

Little MP, Bazyka D, Bouffler SD, Harrison JD, Cardis E, Cucinotta FA, Kreuzer M, Laurent O, Tapio S, Wakeford R, Zablotska L, Lipshultz SE (2012) Estimating risk of circulatory disease: Little et al. respond. Environ Health Perspect 120:453–454.

Lukkes JL, Watt MJ, Lowry CA, Forster GL (2009) Consequences of post-weaning social isolation on anxiety behavior and related neural circuits in rodents. Front Behav Neurosci 3:1–12.

McArdle K, Harrison T, Harrison D (2013) Does a nurturing approach that uses an outdoor play environment build resilience in children from a challenging background? J Adventure Educ Outdoor Learn 13:238–254.

Nijhof SL, Vinkers CH, van Geelen SM, Duijff SN, Achterberg EJM, van der Net J, Veltkamp RC, Grootenhuis MA, van de Putte EM, Hillegers MHJ, van der Brug AW, Wierenga CJ, Benders MJNL, Engels RCME, van der Ent CK, Vanderschuren LJMJ, Lesscher HMB (2018) Healthy play, better coping: The importance of play for the development of children in health and disease. Neurosci Biobehav Rev 95:421–429 Available at: 10.1016/j.neubiorev.2018.09.024.

Pellis SM, Pellis V (2009) The playful brain: venturing to the limits of neuroscience. London, UK: Oneworld Publications.

Pellis SM, Pellis VC, Ham JR, Stark RA (2023) Play fighting and the development of the social brain: The rat’s tale. Neurosci Biobehav Rev 145:105037 Available at: 10.1016/j.neubiorev.2023.105037.

Sampedro-Piquero P, Zancada-Menendez C, Begega A (2015) Housing condition-related changes involved in reversal learning and its c-Fos associated activity in the prefrontal cortex. Neuroscience 307:14–25 Available at: 10.1016/j.neuroscience.2015.08.038.

Sandseter EBH (2007) Categorising risky play—how can we identify risk-taking in children’s play? Eur Early Child Educ Res J 15:237–252.

Sandseter EBH, Kennair LEO (2011) Children’s risky play from an evolutionary perspective: The Anti-phobic effects of thrilling experiences. Evol Psychol 9:257–284.

Schneider P, Bindila L, Schmahl C, Bohus M, Meyer-Lindenberg A, Lutz B, Spanagel R, Schneider M (2016) Adverse social experiences in adolescent rats result in enduring effects on social competence, pain sensitivity and endocannabinoid signaling. Front Behav Neurosci 10:1–16.

Scholl JL, Afzal A, Fox LC, Watt MJ, Forster GL (2019) Sex differences in anxiety-like behaviors in rats. Physiol Behav 211:112670 Available at: 10.1016/j.physbeh.2019.112670.

Sgro M, Mychasiuk R (2020) Playful genes: what do we know about the epigenetics of play behaviour? Int J Play 9:25–38.

Shanazz K, Dixon-Melvin R, Nalloor R, Thumar R, Vazdarjanova AI (2022) Sex Differences In Avoidance Extinction After Contextual Fear Conditioning: Anxioescapic Behavior In Female Rats. Neuroscience 497:146–156 Available at: 10.1016/j.neuroscience.2022.06.031.

Simpson J, Kelly JP (2011) The impact of environmental enrichment in laboratory rats-Behavioural and neurochemical aspects. Behav Brain Res 222:246–264 Available at: 10.1016/j.bbr.2011.04.002.

Sparling JE, Baker SL, Bielajew C (2018) Effects of combined pre- and post-natal enrichment on anxiety-like, social, and cognitive behaviours in juvenile and adult rat offspring. Behav Brain Res 353:40–50 Available at: 10.1016/j.bbr.2018.06.033.

Špinka M, Newberry RC, Bekoff M (2001) Mammalian play: Training for the unexpected. Q Rev Biol 76:141– 168.

Svoboda J, Telenský P, Blahna K, Bureš J, Stuchlík A (2012) Comparison of male and female rats in avoidance of a moving object: More thigmotaxis, hypolocomotion and fear-like reactions in females. Physiol Res 61:659–663.

Tremblay MS (2011) Systematic review of sedentary behaviour and health indicators in school-aged children and youth. Int J Behav Nutr Phys Act:1–22.

Tremblay MS, Gray C, Babcock S, Barnes J, Bradstreet CC, Carr D, Chabot G, Choquette L, Chorney D, Collyer C, Herrington S, Janson K, Janssen I, Larouche R, Pickett W, Power M, Sandseter EBH, Simon B, Brussoni M (2015) Position statement on active outdoor play. Int J Environ Res Public Health 12:6475–6505.

Valentine G, McKendrick J (1997) Children’s outdoor play: Exploring parental concerns about children’s safety and the changing nature of childhood. Geoforum 28:219–235.

Van Den Berg CL, Pijlman FTA, Koning HAM, Diergaarde L, Van Ree JM, Spruijt BM (1999) Isolation changes the incentive value of sucrose and social behaviour in juvenile and adult rats. Behav Brain Res 106:133–142.

Vanderschuren LJMJ, Trezza V (2014) What the laboratory rat has taught us about social play behavior: role in behavioral development and neural mechanisms. Curr Top Behav Neurosci 16:189–212.

Verharen JPH, Ouden HEM den, Adan RAH, Vanderschuren LJMJ (2020) Modulation of value-based decision making behavior by subregions of the rat prefrontal cortex. Psychopharmacology (Berl):doi: 10.1007/s00213-020-05454-7.

Verharen JPH, Van Den Heuvel MW, Luijendijk M, Vanderschuren, Adan RAH (2019) Corticolimbic mechanisms of behavioral inhibition under threat of punishment. J Neurosci 39:4353–4364.

Von Frijtag JC, Schot M, Van Den Bos R, Spruijt BM (2002) Individual housing during the play period results in changed responses to and consequences of a psychosocial stress situation in rats. Dev Psychobiol 41:58– 69.

Wall VL, Fischer EK, Bland ST (2012) Isolation rearing attenuates social interaction-induced expression of immediate early gene protein products in the medial prefrontal cortex of male and female rats. Physiol Behav 107:440–450 Available at: 10.1016/j.physbeh.2012.09.002.

Weir LA, Etelson D, Brand DA (2006) Parents’ perceptions of neighborhood safety and children’s physical activity. Prev Med (Baltim) 43:212–217.

Whitaker LR, Degoulet M, Morikawa H (2013) Social Deprivation Enhances VTA Synaptic Plasticity and Drug-Induced Contextual Learning. Neuron 77:335–345 Available at: 10.1016/j.neuron.2012.11.022.

Wright IK, Upton N, Marsden CA (1991) Resocialisation of isolation-reared rats does not alter their anxiogenic profile on the elevated X-maze model of anxiety. Physiol Behav 50:1129–1132.

Yokota S, Suzuki Y, Hamami K, Harada A, Komai S (2017) Sex differences in avoidance behavior after perceiving potential risk in mice. Behav Brain Funct 13:1–10.

Zeleznikow-Johnston A, Burrows EL, Renoir T, Hannan AJ (2017) Environmental enrichment enhances cognitive flexibility in C57BL/6 mice on a touchscreen reversal learning task. Neuropharmacology 117:219–226 Available at: 10.1016/j.neuropharm.2017.02.009.

